# Methodological Challenges in Translational Drug Response Modeling in Cancer

**DOI:** 10.1101/731836

**Authors:** Lisa-Katrin Schätzle, Ali Hadizadeh Esfahani, Andreas Schuppert

**Affiliations:** Joint Research Center for Computational Biomedicine (JRC-COMBINE), RWTH Aachen University, Aachen, Germany; Aachen Institute for Advanced Study in Computational Engineering Science (AICES), RWTH Aachen University, Aachen, Germany

## Abstract

Translational models directly relating drug response-specific processes observed *in vitro* to their *in vivo* role in cancer patients constitute a crucial part of the development of personalized medication. Unfortunately, ongoing research is often confined by the irreproducibility of the results in other contexts. While the inconsistency of pharmacological data has received great attention recently, the computational aspect of this crisis still deserves closer examination. Notably, studies often focus only on isolated model characteristics instead of examining the overall workflow and the interplay of individual model components. Here, we present a systematic investigation of translational models using the R-package FORESEE. Our findings confirm that with the current exploitation of the available data and the prevailing trend of optimizing methods to only one specific use case, modeling solutions will continue to suffer from non-transferability. Instead, the conduct of developing translational approaches urgently needs to change to retrieve clinical relevance in the future.

## Introduction

Within the context of a permanently growing interest in precision medicine over the last years, where therapies are intended to be tailored to specific characteristics of individual patients, the study of drug sensitivity prediction for complex diseases, such as cancer, has experienced a tremendous boost (De Niz et al., 2016). The availability of both the computational power to work with complex algorithms and large-scale pharmacological datasets gave rise to various drug sensitivity studies. Since *in vitro* experiments are easily standardizable, readily quantifiable and feasible for high-throughput settings, cell lines moved up to become convenient test specimens to explore the characteristics of diverse diseases and the mechanisms behind drug action.

Thus, cell lines were not only extensively characterized by means of their molecular profile, such as mutational status, gene expression, proteomics, copy number variation or methylation, but also based on their responses to broad panels of drugs (Garnett et al., 2012; Shoemaker, 2006; Lamb et al., 2006; Daemen et al., 2013; Barretina et al., 2012; Cancer Cell Line Encyclopedia Consortium and Genomics of Drug Sensitivity in Cancer Consortium., 2015). Consequently, a variety of computational approaches were developed in order to connect the disease-specific molecular profiles and drug responses (Menden et al., 2013; Jang et al., 2014; Smirnov et al., 2015; Luna et al., 2016; Cokelaer et al., 2017). In this context, however, it became apparent that preclinical biomedical research was facing major issues of lacking consistency between studies and exhibiting irreproducibility of experimental findings (Prinz et al., 2011; Baker, 2015). More so, a large percentage of drugs were observed to be failing in the second phase of clinical trials, even though the preclinical investigation had shown promising efficacy. As a result, more and more studies were also targeted at this so-called ”irreproducibility crisis”, demanding standards for study design, measurements, experimental controls, protocols, data management and statistical analyses (Haibe-Kains et al., 2013; Jarvis and Williams, 2016).

There have been enormous efforts to address the data-related aspects of this crisis, by for example introducing the FAIR principles - Findability, Accessibility, Interoperability, and Reusability - to ensure transparency, reproducibility, and reusability of scientific data (Wilkinson et al., 2016). The computational aspect however, still leaves room for improvement. In order to keep an overview of the versatile field of drug sensitivity prediction, efforts were made to systematically compare existing approaches, for example in collaborative projects, such as the DREAM challenges by Costello et al. (2014). Albeit these efforts are great in comparing and improving distinct components of a model, such as batch effect correction methods (Lazar et al., 2012), feature selection methods (Saeys et al., 2007) or regression algorithms (De Niz et al., 2016), they often lack the consideration of the complete modeling workflow and the interplay of the individual pipeline components. Moreover, published methods are inclined to be biased towards the authors’ fields of expertise, which hinders a fair and objective benchmarking of existing methods.

On a different note, even though studies have shown that cell lines reflect many important aspects of human *in vivo* biology, they also exhibit significant differences. This is especially evident regarding the absence of an immune system and a micro-environment of tumor cells in cell cultures, making the direct translation of preclinical models into a clinically relevant context rather difficult (Gillet et al., 2013). Translational models that relate *in vitro* and *in vivo* mechanisms already in the model building process (Mourragui et al., 2019), help to tackle the discrepancy between abstract training of cell line-based models and their clinical application. One prominent solution is to address the genomic differences between *in vitro* and *in vivo* data via batch-effect removal concepts (Geeleher et al., 2014). However, corresponding to the challenges that are opposing pure cell line models, the resulting translational models are not robust across different patient data sets. Moreover, the processes that are relevant to the fitting often remain invisible.

Accordingly, in this study, we intend to systematically investigate the workflow of translational models, using our R-package FORESEE (Turnhoff and Hadizadeh Esfahani et al., 2019) to optimize with respect to all relevant model components and their interplay. By including publicly available approaches and running the different modeling pipelines automatically, we reduce the bias introduced by the users’ heterogeneous experiences in those methods. Furthermore, we hope to inherently attain clinical relevance of our models by directly evaluating the resulting models on patient tumor data. Along with the efforts of identifying promising settings for translational drug response models, we address common pitfalls in this context and work out guidelines that we hope will advance the development and application of more elaborate models in the future, which will ultimately offer valuable insights into applicable precision medicine.

## Results

### A systematic scan of the model space reveals models of outstanding performance

In order to systematically scan the space of translational models, the FORESEE pipeline was applied to different combinations of cell lineand patient data sets, combining five methods of cell response preprocessing with seven approaches of homogenization and batch effect correction, four kinds of feature selection, five ways of feature preprocessing and seven model algorithms. The scan of these 3920 different modeling pipelines revealed models of very high performance. Especially for the breast cancer patient data set GSE6434 (Chang et al., 2005), multiple good model pipelines could be found, with 128 of them yielding an AUC of ROC ¿ 0.81, which is the reported performance of the precedent-setting translational ridge regression model by Geeleher et al. (2014). The best model pipeline, implementing a kmeans binarization of the reported IC50 values, a remove unwanted variation (RUV) homogenization between the cell line- and patient data set, a landmark gene filter, no further feature preprocessing of the remaining gene expression features, and an elastic net or a lasso regression algorithm, which both accomplished a performance of AUC of ROC = 0.99 (Panel A in Figure 1), produced a near to perfect separation between responders and non-responders of the treatment (Panel B in Figure 1). Still, the 3920 different modeling pipelines manifested a high heterogeneity with a wide performance distribution (Panel C in Figure 1) and a median AUC of ROC of 0.55. As portrayed in Figure S1, each of the other patient data sets shows a similarly wide-spread performance distribution for the 3920 models tested with FORESEE. Yet, models of very high performance can be found for each of them, the best model settings being listed in Table 1.

**Figure 1:**
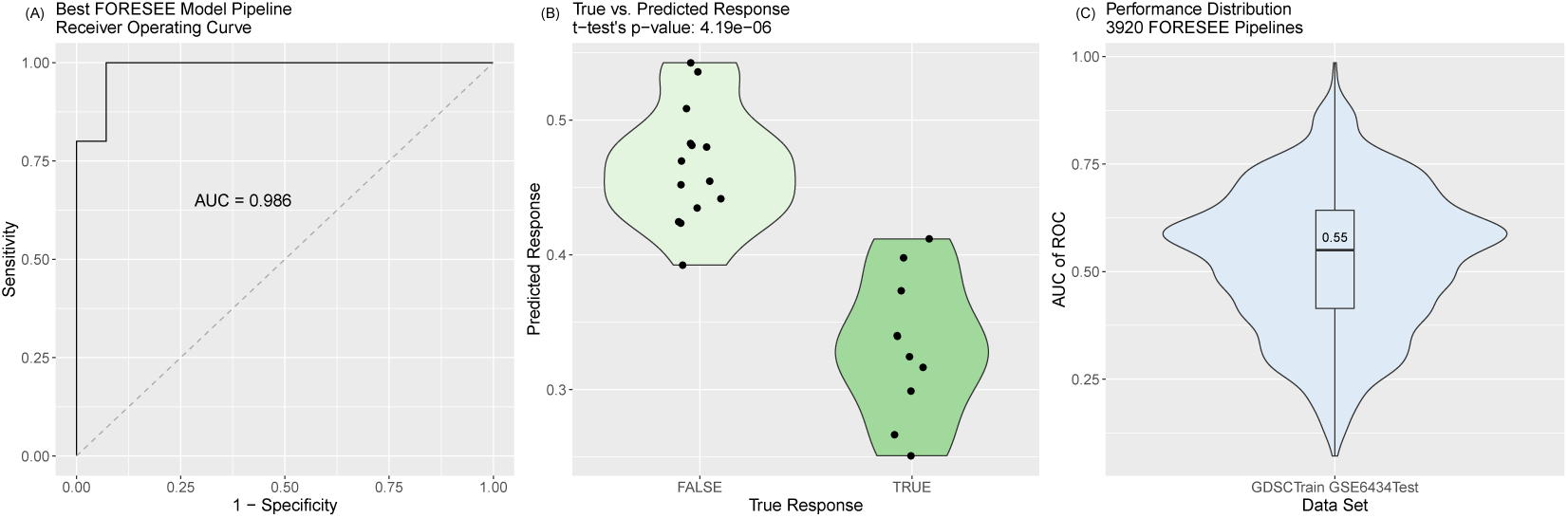
Portrayal of translational models that used the FORESEE package to train models on GDSC cell line data and subsequently predicted GSE6434 patient drug response. Of a total of 3920 modeling pipelines, the best modeling pipeline had the following settings: drug: Docetaxel, cell response type: ln(IC_50_), cell response transformation: binarization with kmeans, sample selection: all, duplication handling: remove all duplicates, homogenization: remove unwanted variation, feature selection: landmark genes, feature preprocessing: none, black box algorithm: elastic net (while lasso would have yielded the same performance). (A) The receiver operating curve of the best model reveals an AUC of 0.986. (B) The comparison of the true responders and non-responders and their separation obtained from the best FORESEE model shows an almost perfect distinction, with a p-value of a ttest of 4.19e-6. (C) The performance distribution of all 3920 model pipelines reveals a median AUC of ROC of 0.55.

**Table 1:**
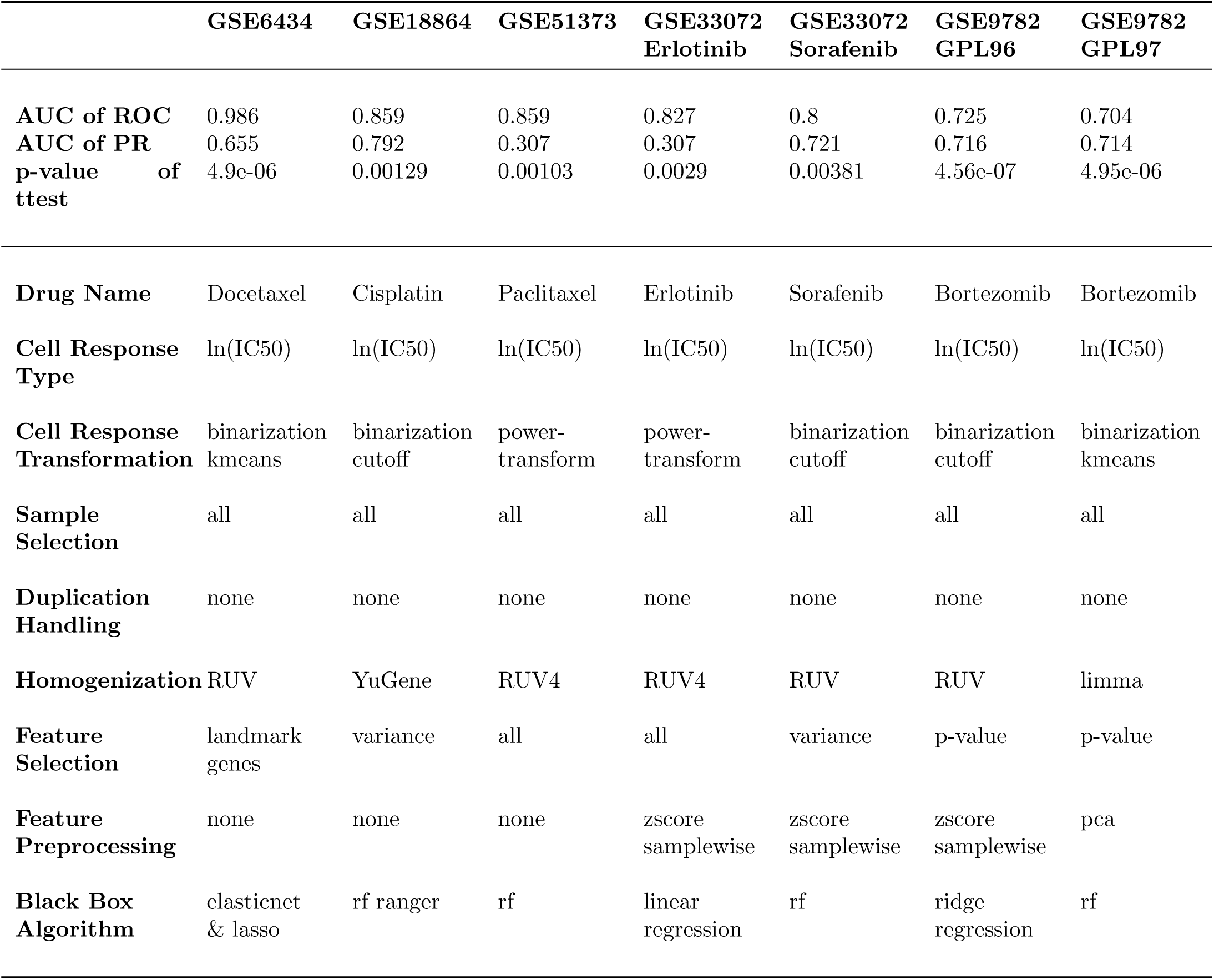
Model Performance and Pipeline Settings of the Best Model for each Patient Data Set.

### Model performance is inconsistent among differing patient data sets

#### A perfect model for a specific data set is insignificant for other use cases

From the comparison of the settings of the best model in each of the cell line - patient data set scenarios in Table 1 it becomes apparent that the best model pipeline settings vary strongly between the different patient data sets. One specific modeling pipeline that yields the best performance in all of the data sets does not exist. In fact, a model that yields almost perfect prediction performance in one of the data sets has a merely mediocre performance for other data sets, as presented in Figure 2, where Panel A shows the AUC of ROC of the best model of each of the analyzed patient data sets Table 1) and the performances of the same models, when they are used to predict the other patient data sets. Panel B of Figure 2 depicts the same information, but in ranks rather than absolute numbers. For example, the best modeling pipeline that was found for modeling the Docetaxel response of GSE6434 patients - here the elastic net regression model that was trained on kmeans binarized IC50 values of GDSC cell lines and RUV homogenized gene expression values of the set of landmark genes without any further preprocessing of the features yields an AUC of ROC = 0.99 in this setting, while for the Cisplatin response of EGEOD18864 patients the same model only generates an AUC of ROC = 0.55, which is ranked 1500th among all 3920 pipelines.

**Figure 2:**
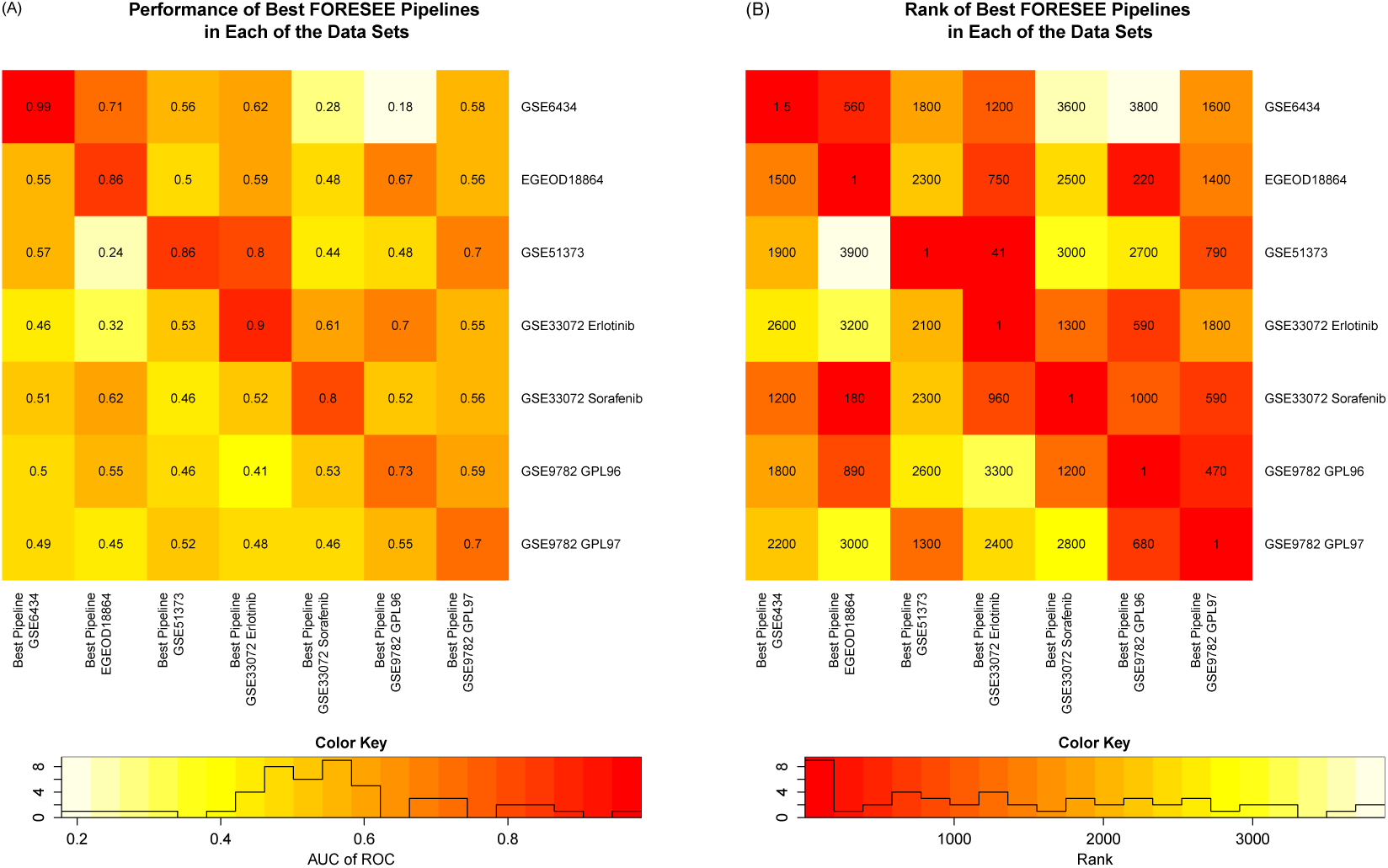
Heatmaps of the performance of the best modeling pipeline of each patient data set (out of 3,920 FORESEE pipelines) in each of the other patient data sets. In (A), the color depicts the AUC of ROC of the respective pipelines, while in (B), the color represents the rank of the modeling pipeline among all 3,920 pipelines that were trained for a specific data set. More details of the modeling pipelines are listed in Table 1. *The rank of the best pipeline of the GSE6434 data set is 1.5, as the modeling pipeline that used lasso instead of elastic net regression for the black box algorithm yielded the exact same performance.

#### Model setting superiority shows generally poor reproducibility among data sets

From our systematic scan of model space, it becomes clear that the problem of reproducibility does not only occur for the very best pipeline in each specific data set, as shown in the section A perfect model for a specific data set is insignificant for other use cases. This problem manifests for almost all modeling pipelines, independent of the absolute value of their performances.

Figure 3 summarizes the performances of 3920 FORESEE modeling pipelines in seven different patient data sets. Instead of seeing clusters of columns, where certain groups of modeling pipelines show identically good or bad performances regardless of the data sets they are applied to, no obvious formation of larger clusters can be identified. This implies that it is impossible to reliably determine if a certain modeling pipeline will yield a good (or bad) prediction performance in another data set, even though the performance in one of the data sets is known. This perfectly mirrors the issue of existing approaches, where the good performance of a model optimized for a specific use case cannot be verified in another situation.

**Figure 3:**
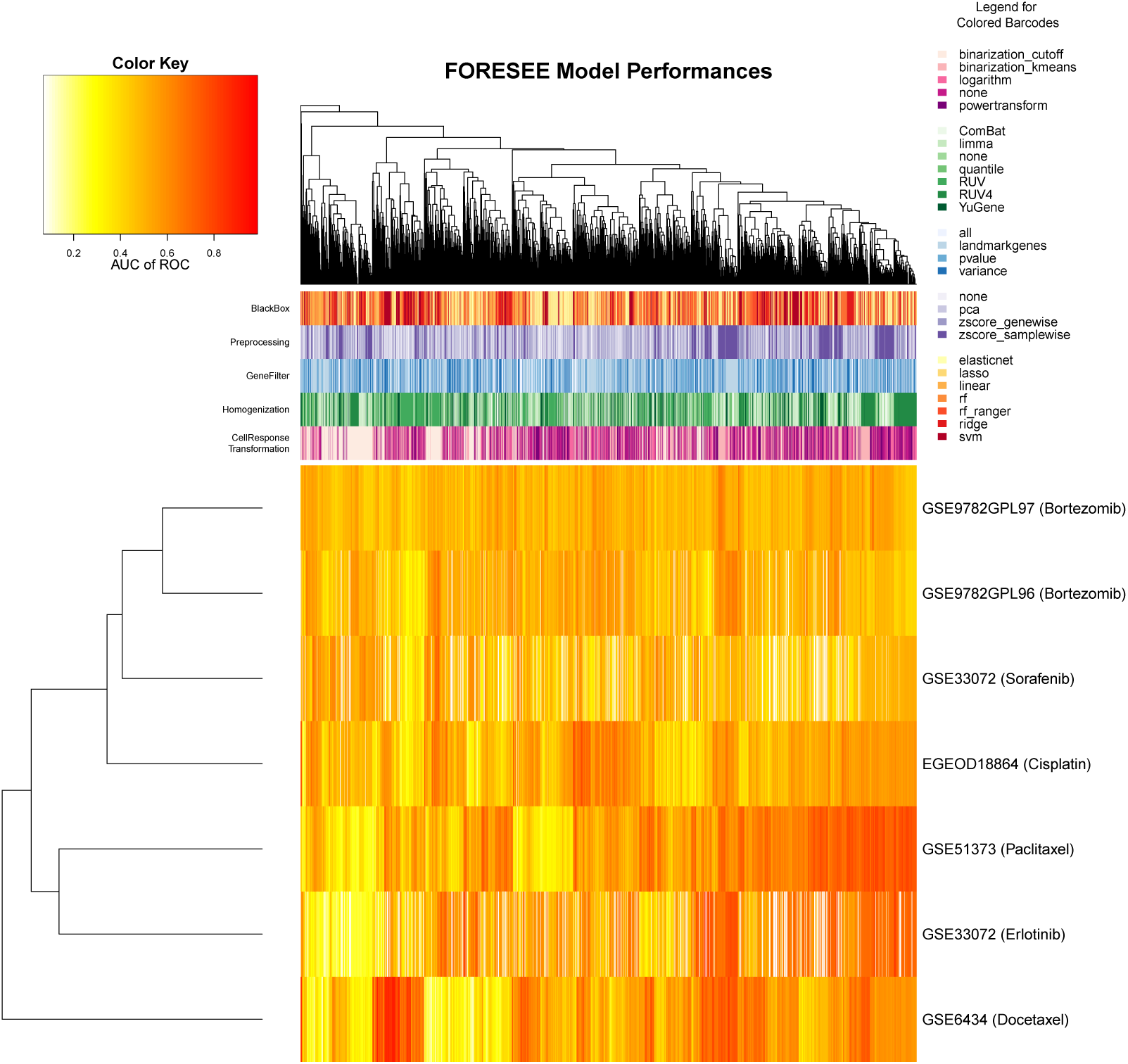
Heatmap of the performance of 3920 translational FORESEE modeling pipelines trained on GDSC cell line data and tested in seven different patient data sets: GSE6434, EGEOD18864, GSE51373, GSE33072 Erlotinib cohort, GSE33072 Sorafenib cohort, GSE9782 GLP96 cohort and GSE9782 GLP97 cohort. The color bars on top of the heatmap represent the individual specifications of each pipeline in the categories CellResponseTransformation, Homogenization, GeneFilter, Preprocessing and Blackbox. The color scheme of the heatmap represents the performance of each pipeline measured as AUC of ROC. The clustering is based on euclidean distance.

Supplementary Figure S2 reinforces this conclusion, as it portrays the Pearson correlation of the AUC of ROC performance values of 3920 modeling pipelines in different patient data sets. Not only are the absolute correlation values very low, with a maximum correlation of 0.49 between GSE51373 and the Erlotinib cohort of GSE33072, but also some of the patient data set pairs exhibit a negative correlation. Especially the correlation between the GPL96 and the GPL97 cohort of the GSE9782 data sets, which comprises the gene expression data of the exact same patients only measured with two different array technologies, exposes a surprisingly low value of 0.41.

In order to put the poor reproducibility into perspective and examine the problem in a broader context, Supplementary Figure S3 features the conditions of the underlying data sets that were used in the presented analyses. After all, there are immediate factors that affect the model performance that do not originate from model specifications.

First, even though all of the models were trained on cell line data from GDSC, the training data sets differed enormously. This is owed to the fact that the drugs included in the GDSC data base are not screened on all of the available cell lines, but mostly on smaller subsets. Consequently, the training data sets for the different translational scenarios in this paper varied in sample size and in the distribution of the cell lines’ tissues of origin depending on the drug that was modelled (Panel C).

Second, the features that were available for training the translational model varied significantly. Both, the micro arrays that were used in the generation of the patient gene expression data and the gene IDs that were originally chosen to define the features, have an impact on the resulting overlapping feature list for each distinctive pair of cell line and patient data set. After converting the reported gene names into Entrez Gene IDs (Maglott et al., 2010) with biomaRt (Durinck et al., 2005, 2009) and removing all duplicates, the feature set size varied from 4786 genes in translational models for the GSE9782 GPL97 data set to 15703 genes in translational models for the GSE33072 Erlotinib and Sorafenib data sets before any feature selection method was applied. As a consequence, all of the gene filter methods *none*, *variance* and *pvalue* produced feature sets that could differ in size by a factor of 3, which makes the comparison of these methods in two different data sets unbalanced.

Third, while the training sets’ distributions of gene expression values was highly similar in all seven settings (Panel A) - which is expected, since all training sets are subsets of the same cell line data base GDSC by Garnett et al. (2012) - the distribution of gene expression differed enormously between the different patient data sets (Panel B). For this very reason, homogenization of the training and test data set pairs is a highly important element of the modeling pipeline. The common homogenization methods should be well prepared for these differences, whereas the option *none*, which simply skips any type of homogenization between the training and test set, can result in very different conditions for different patient data sets. Consequently, it is no surprise that some of the pipelines do not show a high correlation among the different patient data sets. Taking into account these effects of the data constitution, it becomes accessible that the correlation of the ensemble of 3920 pipelines is not in a perfect range. Still, they do not justify the large extent of the bad correlation that was attained in this study and the lack of transferability of these approaches in general.

### Certain model settings can improve translational model performances

As shown in the previous analyses, there is no modeling pipeline that yields outstanding prediction performance for all data sets that were investigated. Yet, it is possible to identify settings that help to generate more promising models.

#### An enrichment analysis reveals that simpler models yield higher performances

A closer investigation of which pipeline settings were enriched among the best model performances for each individual data set provided important insights into promising modeling choices. Table 2 summarizes the significantly enriched model settings in the best 5% of all 3920 FORESEE pipelines in each of the patient data sets partitioned into the five varying pipeline categories: cell response transformation, homogenization, feature selection, feature preprocessing and black box algorithm. Similar to the comparison of the best FORESEE modeling pipeline in each patient data set in Table 1, the significantly enriched model settings vary among the different data sets. Yet, some regularities can be observed for the best performing modeling pipelines:

*•* Binarization of the cell response, using either a cutoff value or the k-means algorithm, is significantly enriched in most of the data sets.
*•* Using a variant of the Remove Unwanted Variation concept for the homogenization of the data sets yields better performances in the majority of data sets.
*•* Reducing the features to genes of the landmark gene list of the LINCS consortium during feature selection is beneficial in four of the seven data sets.
*•* Reducing the dimensionality of the input data by applying principal component analysis to the gene expression values is advantageous in four of the seven data sets.
*•* There is no clearly superior black box algorithm.

**Table 2:** Enrichment of FORESEE Modeling Pipeline Settings in the Best 5% of all Models for Each Patient Data Set.

|  | GSE6434 | GSE18864 | GSE51373 | GSE33072<br>Erlotinib | GSE33072<br>Sorafenib | GSE9782<br>GPL96 | GSE9782<br>GPL97 |
| --- | --- | --- | --- | --- | --- | --- | --- |
| **Cell Response Transformation** | binarization<br>kmeans<br>$p = 3.7e-10$ | binarization<br>cutoff<br>$p = 3.61e-3$ | power-<br>transform<br>$p = 8.96e-3$,<br>none<br>$p = 4.06e-22$ | | binarization<br>cutoff<br>$p = 3.12e-40$ | binarization<br>cutoff<br>$p = 6.61e-3$,<br>logarithm<br>$p = 7.98e-3$ | binarization<br>kmeans<br>$p = 2.65e-9$ |
| **Homogenization** | RUV<br>$p = 3.76e-9$ | YuGene<br>$p = 4.87e-7$ | RUV4<br>$p = 8.67e-42$ | RUV<br>$p = 2.08e-3$ | RUV<br>$p = 2.68e-4$ | RUV<br>$p = 5.89e-3$ | RUV4<br>$p = 5.3e-19$ |
| **Feature Selection** | landmark<br>genes<br>$p = 2.21e-4$ | landmark<br>genes<br>$p = 2.73e-19$ | p-value<br>$p = 3.26e-3$ | landmark<br>genes<br>$p = 3.06e-13$ | | landmark<br>genes<br>$p = 4.89e-16$ | |
| **Feature Preprocessing** | none<br>$p = 1.21e-4$,<br>zscore<br>samplewise<br>$p = 6.44e-5$ | | pca<br>$p = 3.37e-5$ | pca<br>$p = 8.93e-38$ | none<br>$p = 1.43e-6$ | pca<br>$p = 2.37e-25$ | zscore<br>samplewise<br>$p = 3.96e-4$,<br>pca<br>$p = 5.37e-11$ |
| **Black Box Algorithm** | svm<br>$p = 2.29e-18$ | rf<br>$p = 8.17e-20$ | linear<br>regression<br>$p = 4.18e-13$,<br>ridge<br>regression<br>$p = 1.15e-26$ | linear<br>regression<br>$p = 2.03e-4$ | linear<br>regression<br>$p = 3.73e-7$,<br>ridge<br>regression<br>$p = 2.97e-5$ | svm<br>$p = 1.17e-6$ | rf ranger<br>$p = 1.41e-4$ |

Taking into account all categories together, it becomes explicit that settings that aim to simplify the model, such as binarizing the output data or reducing the dimensionality of the input data, tend to be significantly enriched in well performing models and are therefore good principles to aim for.

#### Only few model settings reliably obtain a reasonable performance in multiple data sets

In the context of finding low-risk guidelines for translational modeling, it can be worthwhile to expose rules for a model design that is to a certain extent optimized with respect to a reliably good performance in different model applications rather than one specific use case or data set only, even though the absolute performance value does not seem outstanding when considered alone. In this regard, it is possible to find models that successfully predict 5 out of 7 patient data sets with an AUC of ROC of 0.65 or higher. Intriguingly, there are only three models that fulfill this condition. They are summarized in Table 3. In alignment with the results of the enrichment analysis, key characteristics of a robust model are binarizing the cell line IC50 values, homogenizing the two data sets with a RUV method, and using principal component analysis for feature preprocessing. In contrast to the results of the enrichment analysis however, robust models comprise penalized regression models, such as lasso and elastic net, as black box algorithms, which are not significantly better performing in any of the patient data sets when they are considered individually. Notably, both the GSE18864 data set and the GSE33072 Sorafenib cohort fall short of the cutoff for all three pipelines listed in Table 3, exhibiting near to random performance, while those pipelines behave similarly for the other data sets, suggesting that the difference might originate from the data sets themselves rather than the pipelines.

**Table 3:** Summary of the three FORESEE modeling pipelines that yield an AUC of ROC of 0.65 or higher in at least 5 out of 7 patient data sets.

|  | **Pipeline 1** | **Pipeline 2** | **Pipeline 3** |
| --- | --- | --- | --- |
| **Cell Response Transformation** | binarization kmeans | binarization kmeans | binarization kmeans |
| **Homogenization** | RUV4 | RUV4 | RUV4 |
| **Gene Filter** | pvalue | pvalue | variance |
| **Preprocessing** | pca | pca | pca |
| **Black Box Algorithm** | elastic net | lasso | lasso |
| **AUC of ROC for GSE6434** | 0.75 | 0.75 | 0.74 |
| **AUC of ROC for GSE18864** | 0.57 | 0.57 | 0.57 |
| **AUC of ROC for GSE51373** | 0.72 | 0.72 | 0.68 |
| **AUC of ROC for GSE33072 Erlotinib** | 0.71 | 0.71 | 0.70 |
| **AUC of ROC for GSE33072 Sorafenib** | 0.50 | 0.51 | 0.51 |
| **AUC of ROC for GSE9782 GPL96** | 0.66 | 0.66 | 0.67 |
| **AUC of ROC for GSE9782 GPL97** | 0.66 | 0.67 | 0.67 |

### Investigation of Randomness

From the exploration of the lack of correlation of model performances in the different data sets and rather weak coherences for reliable model settings in general, it becomes apparent that the translational models of this study are affected by noise. Therefore, we aspired to assess the extent of randomness in the results by two different approaches.

#### The sample size of the patient data sets affects the perception of noisy results

As apparent from the violin plots in Figure 1 and in Supplementary Figure S1, models seem to yield much higher performances for data sets such as GSE6434 with a maximum AUC of ROC of 0.986 than data sets such as the GSE9782 GPL97 cohort with a maximum AUC of ROC of only 0.704. While there is no doubt that noise varies from data set to data set, such that certain ones are better adapted for model training than others, it is still reasonable to investigate the circumstances of this performance disparity in more detail.

One principal difference of the patient data sets is their sample size. Therefore, Figure 4 displays the performance distributions of all 3920 modeling pipelines in the order of increasing sample size of the patient data sets (light blue violin plots). Following the principle that it is a lot more probable to correctly guess the responses of a small sample set than those of a very large sample set at random, it becomes explicit that data sets that exhibit a wide spread of performance values with high reaching peak performances have a significantly smaller sample size (GSE6434: 24 patients, EGEOD18864: 24 patients and GSE51373: 25 patients) than data sets with a narrow performance distribution (GSE9782 GPL96 and GPL97: 169 patients). For the purpose of illustration of this sample size effect, Figure 4 includes distributions that represent artificial AUC of ROC values from the comparison of 10,000 randomly generated binary vectors in the size of the patient data sets compared to the actual responses of the respective patient data sets (dark blue). Moreover, Figure 4 depicts distributions that summarize the performances of the 3,920 FORESEE model pipelines applied to 1,000 versions of each patient object, where the gene labels were randomly permuted (medium blue).

**Figure 4:**
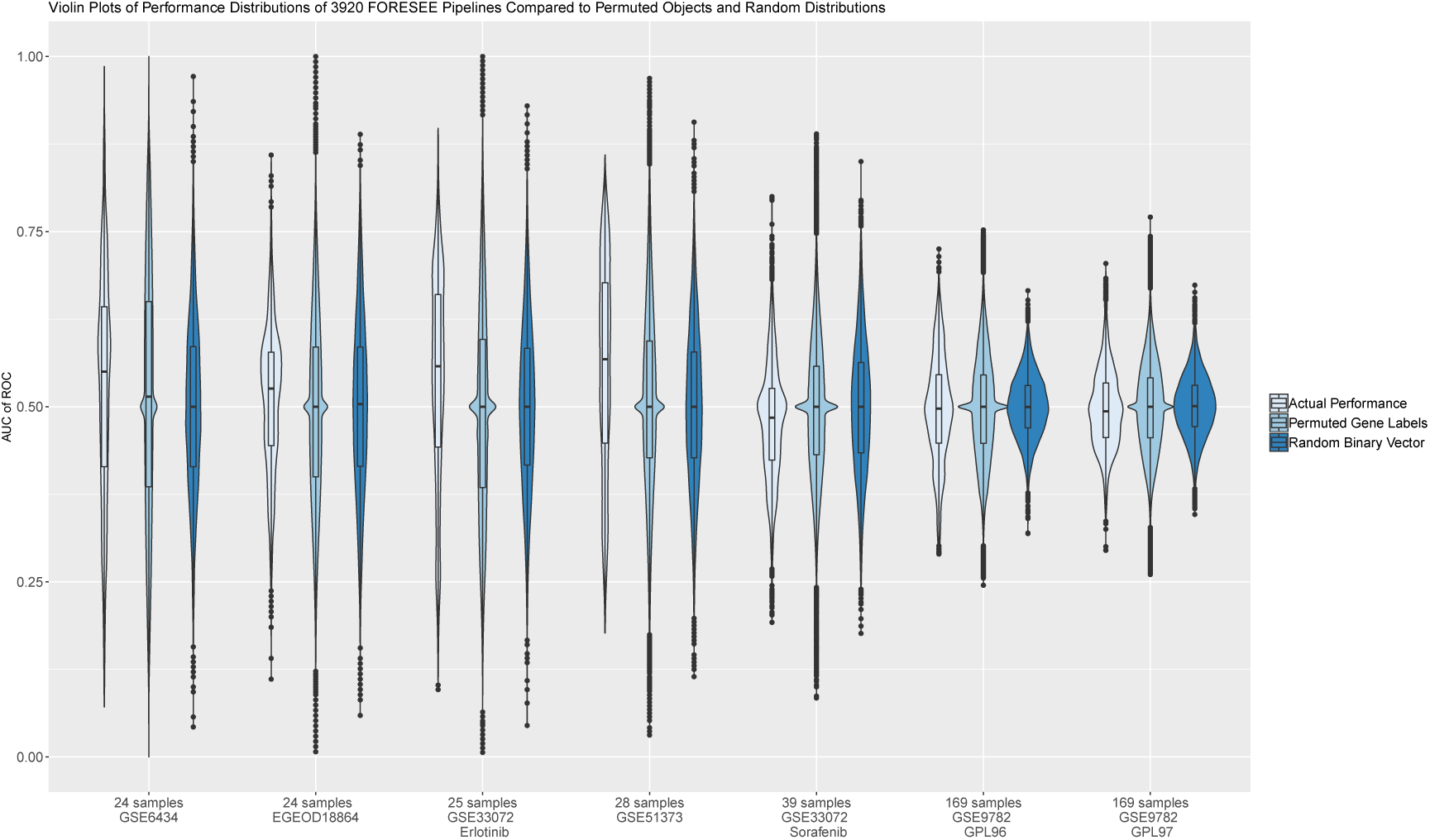
Violin plots of the performances of each of the seven different patient data sets: GSE6434, EGEOD18864, GSE51373, GSE33072 Erlotinib cohort, GSE33072 Sorafenib cohort, GSE9782 GLP96 cohort and GSE9782 GLP97 cohort. The actual performance distributions of the translational models (light blue) are compared to the distributions, where each of the 3,920 translational models was applied to 1,000 patient objects with randomly permuted gene labels (medium blue) and to distributions, where the actual patient response values are compared to 10,000 randomly generated binary vectors to calculate an artificial AUC of ROC measure (dark blue). Data sets are shown in increasing sample size from left to right.

From the direct comparison between the actual performance distributions and the random ones it becomes apparent that certain model findings may be less significant than previously thought. In the cases of GSE9782 GPL96 and GPL97 for instance, even though the best models have comparably low absolute values of AUC of ROC, their performances are much higher than predictable by 10,000 random guesses. Thus, they effectively allow the training of robust models that are not driven by noise. The Sorafenib cohort of the GSE33072 data set and the GSE18864 data on the other hand yield worse results on average when a model is trained to predict the response than when the response is guessed at random, even though the absolute values of the AUC of ROC seem more promising than the ones of GSE9782 GPL96 and GPL97. This is particularly prominent for the Sorafenib cohort of the GSE33072 data set, which exhibits a median AUC of ROC of 0.48 - the lowest among all data sets.

Combined with the finding that the GSE18864 data set and the GSE33072 Sorafenib cohort fall out of line for the robust pipelines listed in Table 3, this indicates that these two data sets are predominantly governed by noise and therefore unsuitable for the training of universally valid clinical models. For GSE18864, this finding is concurrent with previous findings by Geeleher et al. (2014), whose models could not capture variability in clinical response, and by Silver et al. (2010), the authors of the original study, who could not identify a predictive gene signature of Cisplatin response. Similarly, a unified set of biomarker genes that establishes a robust prediction of Sorafenib responses for the GSE33072 data set has not been determined so far, for example by Blumenschein et al. (2013).

Even though the other data sets show a better performance distribution of the 3920 FORESEE models than the GSE18864 and the GSE33072 Sorafenib data sets, it becomes explicit that the fitted models do not outperform the randomized models by far. Especially the large range of the performance distribution of translational models applied to patient data with permuted gene labels (medium blue) demonstrates that an optimized model of very high predictive performance in one application scenario - as listed in this study or as published separately by other groups - does not necessarily produce a model that truly captures the mechanisms that are relevant for the drug response and can therefore not be employed in clinical practice. Instead, the high predictivity can equiprobably stem from random mechanisms that were captured by the model by chance and are only relevant to that specific scenario. In this sense, this analysis demonstrates the importance of considering the possible degree of randomness that could affect modeling results, especially when the sample size of the test population is small.

### Well performing translational models are not necessarily specific to the drug of interest

In an attempt to further analyze the extent of randomness accompanying the performance results of the shown FORESEE modeling pipelines, a drug specificity analysis was conducted. For each of the patient data sets and for each of the 266 drugs contained in the GDSC data base, a set of 15 randomly chosen pipelines, which are summarized in Table 4, was used to train translational models on the GDSC cell line data and predict the respective patient response. Figure 5 depicts the drugs that were used during the training process for each indivitual patient data set ranked according to their mean performance. In none of the cases the drug that is administered to the patients is the best option to pick for training the translational models on the cell line data. While for some of the data sets, namely the Docetaxel treated breast cancer patients of GSE6434 and the Paclitaxel treated ovarian cancer patients of GSE51373, the specific drug that is administered to the patient is in the top range of training drugs, for other data sets, such as the Bortezomib treated multiple myeloma patients of both GSE9782 GPL96 and GPL97, as well as the Cisplatin treated breast cancer patients of GSE18864, the drugs of interest only have mediocre ranks compared to other drugs. Especially the latter examples give the impression that the drug that is used during the training process has only a minor effect on the model performance. Indeed, the fact that so many models reveal good performances, even though the drugs that the models were trained with are not the ones that were predicted in the end, suggests that the internal mechanisms that dominate the translational models for drug action in cancer patients are not specific to the drug of interest, but rather to general mechanisms of drug response. This hypothesis is further supported by the observation that the drugs that are more predictive than the drugs of interest do not share the same mode of action. For example, Docetaxel is a chemotherapeutic drug that binds to microtubules and prevents their disassembly, eventually causing the initiation of apoptosis, while the drug occupying the first rank in predicting GSE6434 patients, Tipifarnib, is a Farnesyl-transferase inhibitor that ultimately prevents the proper functioning of Ras. Likewise, Bortezomib is a proteasome inhibitor, while the drug that is ranked first for GSE99782 GPL97 patients is Quizartinib, a FLT3 kinase inhibitor.

**Figure 5:**
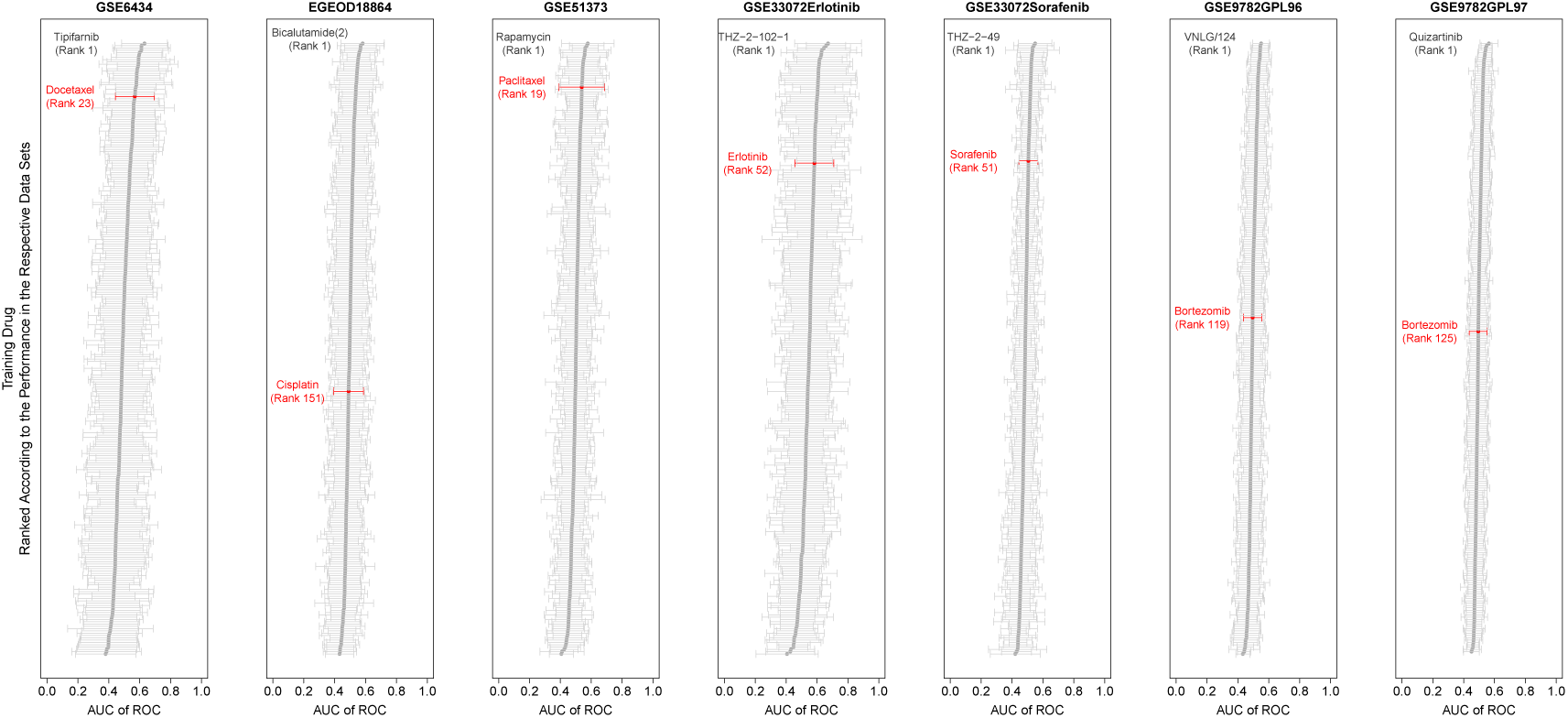
Drug specificity plot of each of the seven different patient data sets: GSE6434, EGEOD18864, GSE51373, GSE33072 Erlotinib cohort, GSE33072 Sorafenib cohort, GSE9782 GLP96 cohort and GSE9782 GLP97 cohort. A set of 15 pipelines, which are summarized in Table 4, were randomly chosen and used to train translational models on the GDSC cell line data with each of the 266 drugs contained in the GDSC data base individually and then tested on each of the patient data sets. For each of the data sets, the drugs are ordered with respect to the mean AUC of ROC of the 15 random pipelines trained with that drug. The red color marks the drug that is actually applied to the patient. The first-ranked drug is additionally indicated in order to facilitate the comparison of the different drugs and their modes of action.

**Table 4:** Pipeline Settings for the Drug Specificity Analysis. 15 random FORESEE pipelines were chosen and then trained on every available drug in the GDSC data base to predict the patient data sets.

| Pipeline | Response Type | Cell Response Transformation | Sample Selector | Duplication Handling | Homogenization | Gene Filter | Preprocessing | Black Box |
| --- | --- | --- | --- | --- | --- | --- | --- | --- |
| **1** | LN_IC50 | binarization cutoff | all | none | ComBat | pvalue | pca | elasticnet |
| **2** | LN_IC50 | binarization cutoff | all | none | limma | variance | zscore samplewise | lasso |
| **3** | LN_IC50 | binarization cutoff | all | none | none | pvalue | zscore samplewise | elasticnet |
| **4** | LN_IC50 | binarization cutoff | all | none | quantile | landmark-genes | zscore genewise | elasticnet |
| **5** | LN_IC50 | binarization cutoff | all | none | RUV | pvalue | zscore samplewise | linear |
| **6** | LN_IC50 | binarization kmeans | all | none | quantile | variance | zscore genewise | lasso |
| **7** | LN_IC50 | logarithm | all | none | ComBat | landmark-genes | zscore genewise | svm |
| **8** | LN_IC50 | logarithm | all | none | RUV | landmark-genes | zscore genewise | lasso |
| **9** | LN_IC50 | logarithm | all | none | RUV4 | landmark-genes | zscore samplewise | ridge |
| **10** | LN_IC50 | logarithm | all | none | RUV4 | pvalue | none | lasso |
| **11** | LN_IC50 | logarithm | all | none | RUV4 | variance | none | svm |
| **12** | LN_IC50 | none | all | none | ComBat | pvalue | none | lasso |
| **13** | LN_IC50 | none | all | none | none | pvalue | zscore genewise | rf_ranger |
| **14** | LN_IC50 | powertransform | all | none | RUV | landmark-genes | pca | svm |
| **15** | LN_IC50 | powertransform | all | none | YuGene | landmark-genes | zscore samplewise | ridge |

## Discussion

The ensemble of investigations of this study proves that a systematic analysis of translational models, not focussing on one specific model element only, but instead considering the whole modeling pipeline and the interplay of preprocessing methods and modeling algorithms, can be beneficial to refine the understanding of modeling coherences and key model characteristics. Tools, such as our R-package FORESEE, have the potential to support this systematic analysis by spanning a space of model parameters and intrinsically optimizing the settings. By considering all combinations of model settings, this optimization indeed spawned superior models for individual application scenarios. This superiority however was proven to not be robust across multiple data sets, therefore mirroring the common problem of failing reproducibility of translational modeling approaches in contemporary literature.

While this seems to resemble the common overfitting problem for models that are trained and tested on one data set only and fail to replicate in a new, unknown one, this finding is different in the sense that during the translational model fitting process, a model is trained on cell line data and it is not until after the training process that the model is applied to the patient data to obtain a prediction. Considered in isolation, this separation of training- and test set should provide for a definite quantification of model performance that is robust with respect to overfitting to the underlying data. The fact that it is not robust closely corresponds with the common failure of reproducibility of known approaches. Strikingly however, the incorporation of the optimization process on the model parameter space seems to introduce an effect that is quite similar to the overfitting problem. With the exception of the minor advantage caused by simplifying translational models by for example binarizing the training output data, reducing the feature list to landmarkgenes, reducing the input dimensionality by PCA or applying penalized regression methods, no model settings or setting combinations could be revealed that reliably promised superior performance for more than one model application. Instead, we could demonstrate that random predictions could in fact yield similar performance values. Moreover, the drug specificity analysis showed that the drug that a translational model is trained with does not have a significant impact on the model performance. Effectively, the training drug does not even have to share the same mode of action with the drug that is predicted to create a good model.

Taken as a whole, this proves that translational models in the fashion of the ones developed in this study are not yet able to explain all processes of drug action in patients. On the contrary, the processes that are dominating translational models are not yet understood, such that random or drug unspecific models seem to be equally good predictors in most of the cases. Yet, the broader concept of using *in vitro* experiments in the preclinical drug development stage to learn drug mechanisms for the subsequent *in vivo* application in patients is the current state-of-the-art in pharmacological research. Even though it seems possible to capture general processes of drug action and survival with translational models, the drug specific processes seem to play a minor part, which could be one reason for the large number of drugs that have to be withdrawn from the market after failing in the clinical phase, even though they had shown promising results in the preclinical phase. This substantiates that more complicated models are needed and that the integration of profound and mechanistic knowledge into the model could be highly beneficial.

Along the way to more insightful models, also other important factors should be considered; some of which emerged within this study. One challenge in the development process of translational models is the extent of utilizable data, which falls in line with the aforementioned investigations concerning the reproducibility crisis of data. Despite the fact that the generation of omics data has experienced a tremendous boost over the last years, the availability of patient data sets that include both molecular data, such as gene expression, and a measure of drug efficacy is limited. Existing patient data sets comprise small sample sizes only, which makes model training with them susceptible to noise, as apparent from the sample size effect analysis. Moreover, these data sets are highly heterogeneous in terms of the used array technologies, the measured drug response type, the chosen gene names and the applied normalization method. The creation of robust models that are easily transferable to other data sets requires large data sets of the same structure and preprocessing, such that they are effortlessly comparable. At the same time, models that are trained on data sets of small sample sizes only, should be thoroughly tested with regard to their randomness and robustness in more than one context before being purported as reliably predictive. To that effect, this study accentuates once more the importance of systematic and unbiased benchmarking of new approaches.

Another important aspect of the exploration of translational models is the reliability of the translational principle as such. With the great amount of noise and randomness in the results of current studies, it should be investigated to what extent cell lines are actually predictive of patients and whether or not other model systems, such as organoids (Fatehullah et al., 2016) or patient-derived xenografts (Dobrolecki et al., 2016) could be of higher similarity with respect to patients and therefore more suitable to understand drug mechanisms.

Lastly, missing predictivity of the translational models shown here could be partially ascribed to the molecular data type that the models were trained on. So far, the investigations of translational models were mainly focused on gene expression data of microarrays owed to the absence of patient data sets that comprise more than one *omics* data type and at least one measure of drug response. The inclusion of RNASeq data and other data types, such as mutational status, copy number variation, methylation and protein expression could significantly improve the predictive performance of models, simply by adding more levels of information. Since not all biomedical mechanisms are fully shown on the gene expression level, other data types could provide valuable insights to generate a detailed picture of the disease and drug mechanisms.

## Methods and Materials

### Data

#### Cell Line Data

For the training of the translational models in this study, both gene expression data and IC_50_ (half maximal inhibitory concentration) values as drug response measure were derived from GDSC data (Garnett et al., 2012) and formatted into *ForeseeCell* objects, which is summarized in Table 5.

**Table 5:** Cell Line Data Sets.

| Data Set | Consortium | Samples | Array | Reference |
| --- | --- | --- | --- | --- |
| GDSC | Genomics of Drug Sensitivity in Cancer | 1,065 | U219 Array* | Garnett et al. (2012) |

\* Affymetrix Human Genome

#### Patient Data

For the testing of the translational models in this study, information of patients with breast cancer (GSE6434 (Chang et al., 2005) and GSE18864 (Silver et al., 2010)), lung cancer (GSE33072 (Byers et al., 2013)), ovarian cancer (GSE51373 (Koti et al., 2013)) and multiple myeloma (GSE9782 (Mulligan et al., 2007)) was organized into *ForeseePatient* objects including gene expression data and one measure of *in vivo* drug efficacy, which is summarized in Table 6. Details about the preparation of the data sets can also be found in the Supplementary File 2 of our FORESEE package publication (Turnhoff and Hadizadeh Esfahani et al., 2019).

**Table 6:** Patient Data Sets.

| GEO Data Set | Cancer Type | Drug | Samples | Responder/ Non-Responder | Array | Reference |
| --- | --- | --- | --- | --- | --- | --- |
| GSE6434 | Breast Cancer | Docetaxel | 24 | 10/14 | U95 Version 2 Array* | Chang et al. (2005) |
| GSE18864 | Breast Cancer | Cisplatin | 24 | 15/9 | U133 plus 2.0 Array* | Silver et al. (2010) |
| GSE51373 | Ovarian Cancer | Paclitaxel | 28 | 16/12 | U133 plus 2.0 Array* | Koti et al. (2013) |
| GSE33072 | Non-Small Cell Lung Cancer | Erlotinib | 25 | 13/12 | 1.0 ST Array* | Byers et al. (2013) |
| GSE33072 | Non-Small Cell Lung Cancer | Sorafenib | 39 | 20/19 | 1.0 ST Array* | Byers et al. (2013) |
| GSE9782 | Multiple Myeloma | Bortezomib | 169 | 85/84 | U133A Array* | Mulligan et al. (2007) |
| GSE9782 | Multiple Myeloma | Bortezomib | 169 | 85/84 | U133B Array* | Mulligan et al. (2007) |

\* Affymetrix Human Genome

### Methods

Analyses of this paper were conducted with the software *R* (R Core Team, 2017).

#### FORESEE

For the systematic comparison of different translational drug response modeling pipelines, the R-package FORESEE by Turnhoff and Hadizadeh Esfahani et al. (2019) was used. FORESEE, which is short for uniFied translatiOnal dRug rESponsE prEdcition platform, partitions the general modeling pipeline into defined functional elements in order to enable the user to thoroughly investigate the impact of each of them on the model performance. The FORESEE modeling routine comprises two major functions: the ForeseeTrain loop, which uses cell line data to train a translational model, and the ForeseeTest loop, which applies the learned model to new patient data and assesses its performance. During training, the cell response data is transformed to serve as model output, while molecular data is prepared to be used as model input: before the data is fed into a black box model, training samples are selected, duplicated features are removed, cell lineand patient data are homogenized by means of batch effect correction methods, and specific features are selected, transformed and combined. For model testing, the completed model is applied to molecular patient data that needs to be preprocessed in the exact same manner as the cell line data. The predictions can subsequently be compared to the reported patient drug responses to evaluate the model performance.

##### Cell Response Preprocessing

Five different methods to preprocess the cell response data were chosen. Three options kept the cell response values continuous in order to train regression models:

1. The method *none* used the reported cell response values without any preprocessing.
2. The method *logarithm* applied the natural logarithm to the response values. In case of negative drug response values, an offset equal to the negative minimum drug response value plus 1 was added to all response values in order to avoid negative arguments of the logarithm.
3. The method *powertransform* used the R-package car by Fox and Weisberg (2011) to determine an exponent for a subsequent power transformation. Again, an offset equal to the negative minimum drug response value plus 1 was added to all response values, in case of negative drug response values.

Additionally, two different binarization methods were used for classification settings:

4. The method *binarization cutoff* used the package bootnet by Epskamp et al. (2018) to split the drug response values at the median into two classes.
5. The method *binarization kmeans* used the package Binarize by Mundus et al. (2017) to define two classes of responders using kmeans clustering.

##### Patient Response Preprocessing

While the patient data sets GSE6434 and GSE51373 already contained binary response annotations, all other patient data sets had to be binarized before usage. The GSE33072 data with Erlotinib and Sorafenib response were binarized by splitting the reported progression free survival time in months at the median. For the patient data set GSE18864, patients, where the clinical response had been categorized as clinical complete response (cCR) or clinical partial response (cPR), were manually classified as responders, whereas patients, where the clinical response had been categorized as stable disease (SD) or progressive disease (PD), were classified as non-responders. Similarly, the responses to Bortezomib of the GSE9782 data set were manually classified into responders, if the response had been reported as complete response (CR), partial response (PR) or minimal response (MR), and non-responders, if no change (NC) or progressive disease (PD) had been reported.

##### Sample Selection

Previous studies have shown an increased predictivity of translational drug response models whose training sets do not only include cell line samples from the same tissue of origin as the tumour, but also cell lines originating from diverse other tissues (Geeleher et al., 2014). Therefore, all available cell line samples from various tissues were selected to train the models in this study.

##### Duplication Handling

In order to provide unique features and avoid any mismatches, all gene names that occurred more than once were removed in both the training and the test object.

##### Homogenization

For the homogenization of the gene expression data and the batch effect correction between cell line train- and patient test data, seven different methods were chosen:

1. The method *ComBat* from the package sva by Leek et al. (2017) used empirical Bayes frameworks to adjust data for batch effects.
2. The method *limma* by Ritchie et al. (2015) removed covariate effects by fitting a linear model to the impact of the different batches.
3. In order to have a thorough investigation of the usefulness of homogenization methods, the option *none* offered to employ modeling pipelines without any correction.
4. The method *quantile* by Bolstad (2017) homogenized the data of different origins with quantile normalization.
5. The method *RUV4* by Gagnon-Bartsch (2018) homogenized the gene expression data sets with the help of a list of housekeeping genes by Eisenberg and Levanon (2013), which are defined by a constant level of expression across tissues. In a singular value decomposition, these housekeeping genes were considered as negative controls to identify and subsequently remove unwanted variation.
6. The method *RUV* applied a self-implemented function that was inspired by the function *RUV4* by Gagnon-Bartsch (2018) and a function by Geeleher et al. (2014) in order to remove batch effects. The function *princomp()* of the *stats* package of R applied a principal component analysis on the gene expression of the housekeeping genes (Eisenberg and Levanon, 2013) of both data sets and determined the impact of unwanted variation by training a linear regression model with the function *lm()* of the *stats* package of R on the first 10 principal components. The residual gene expression of all genes was considered as homogenized data.
7. The method *YuGene* by Lê Cao et al. (2014) applied a cumulative proportion approach to make the gene expression data sets comparable.

##### Feature Selection

Selecting a smaller subset out of the thousands of genes in microarray data to increase the robustness of biological models is a widely studied topic. In this paper, we compare four simple approaches:

1. The method *landmarkgenes* reduced the features to the list of landmark genes that were determined as being informative to characterize the whole transcriptome by Subramanian et al. (2017)
2. The method *variance* used the function *var()* of the *stats* package of R to calculate the variance of each gene across all samples of the training data set. The 20% least variant genes were removed from the analysis.
3. The method *pvalue* calculated a student’s t-test with the function *t.test()* of the *stats* package of R between the most sensitive and the most resistant samples of the training data set. The 20% genes with the highest p-values were removed from the analysis.
4. The option *all* considered all overlapping genes between the train- and test object without further feature selection.

##### Feature Preprocessing

For the standardization and transformation of the features, four different methods were chosen:

1. The method *zscore samplewise* transformed the raw gene expression (GEX) value of each gene, by subtracting the mean GEX of all genes of one sample and subsequently dividing the resulting value by the standard deviation of the GEX of all genes of that sample.

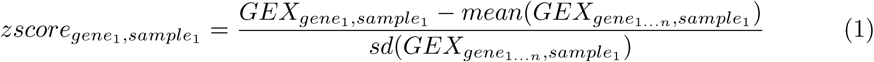
2. The method *zscore genewise* transformed the raw gene expression (GEX) value of each gene, by subtracting the mean GEX of this gene in all samples and subsequently dividing the resulting value by the standard deviation of the GEX of this gene in all samples.

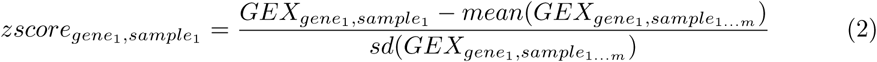
3. The method *pca* used the function *prcomp()* of the *stats* package of R to convert the possibly correlated raw gene expression features into a set of linearly uncorrelated principal component features. The first 10 principal components were chosen as a dimensionality-reduced feature set for the model.
4. The option *none* included the raw gene expression features without any preprocessing to offer a comparison.

##### Feature Combination

All model pipelines in this study were based on gene expression data only, since the patient data sets that were chosen for this study did not contain other molecular features.

##### BlackBox Filtering

For the final step in the model development, seven different algorithms were chosen to train a model that predicts the drug response of cell lines based on baseline gene expression profiles:

1. The method *linear* used the function *lm()* of the *stats* package of R to fit a linear model to the training data.
2. The method *lasso* by Friedman et al. (2010) fitted a generalized linear model with lasso penalty (*α* = 1) and a regularization parameter *λ* determined by a 10-fold crossvalidation.
3. The method *elasticnet* by Friedman et al. (2010); Microsoft and Ooi (2017) fitted a generalized linear model with an elastic net mixing parameter of (*α* = 0.5) and a regularization parameter *λ* determined by a 10-fold crossvalidation
4. The method *rf* by Liaw and Wiener (2002) used a random forest algorithm by Breiman (2001) to fit the training data with 500 trees.
5. The method *rf ranger* by Wright and Ziegler (2017) trained a fast implementation of a random forest model on the training data by training 10,000 trees of unlimited depth.
6. The method *ridge* by Moritz and Cule (2017) fitted a linear ridge regression model to the data, where the ridge parameter was chosen automatically based on the method of Cule and De Iorio (2012))
7. The method *svm* by Meyer et al. (2017) trained a support vector machine with a radial kernel on the training data.

##### Validation

For the investigation of the performance of the different modeling pipelines, the area under the receiver operating curve between the true and the predicted patient classes was calculated using the *pROC* package by Robin et al. (2011).

#### Statistics

##### Significance Testing

In order to investigate whether two distributions were significantly different from each other the function *t.test()* of the *stats* package of R applied a two-sided student’s t-test to the two distributions.

##### Enrichment Analysis

In order to investigate the enrichment of certain model settings in the overall distribution of model performances with a hypergeometric function, the method *phyper()* of the *stats* package of R was applied to the data.

##### Receiver Operating Curves

Receiver operating curves were calculated with the *roc()* function of the *pROC* package by Robin et al. (2011) and plotted with the *ggroc()* function of the *ggplot2* package by Wickham (2016).

##### Correlation

In order to determine the linear correlation of the performance distributions, the method *pearson* measured the Pearson correlation with the function *cor()* of the *stats* package of R.

##### Gene Labeling

Within the context of collecting the different data sets and enforcing them into the same ForeseeObject formats for the *FORESEE* R package (Turnhoff and Hadizadeh Esfahani et al., 2019), the *biomaRt* package by Durinck et al. (2005, 2009) was applied to convert all gene labels into Entrez IDs.

#### Plotting

##### Heatmaps

Heatmaps were plotted with the *heatmap.2()* function of the R package *gplots* by Warnes et al. (2019) or the *heatmap.3() function* by Griffith (2016).

##### Histograms

All histograms were plotted with the *hist()* function of the *graphics* package of R.

##### Violin Plots

Violin plots were created using the *ggplot() + geom violin()* function of the *ggplot2* package by Wickham (2016).

##### Correlation

The correlation plot was created using the *corrplot.mixed()* function of the *corrplot* package by Wei and Simko (2017).

## Supplementary Material

**Figure S1:**
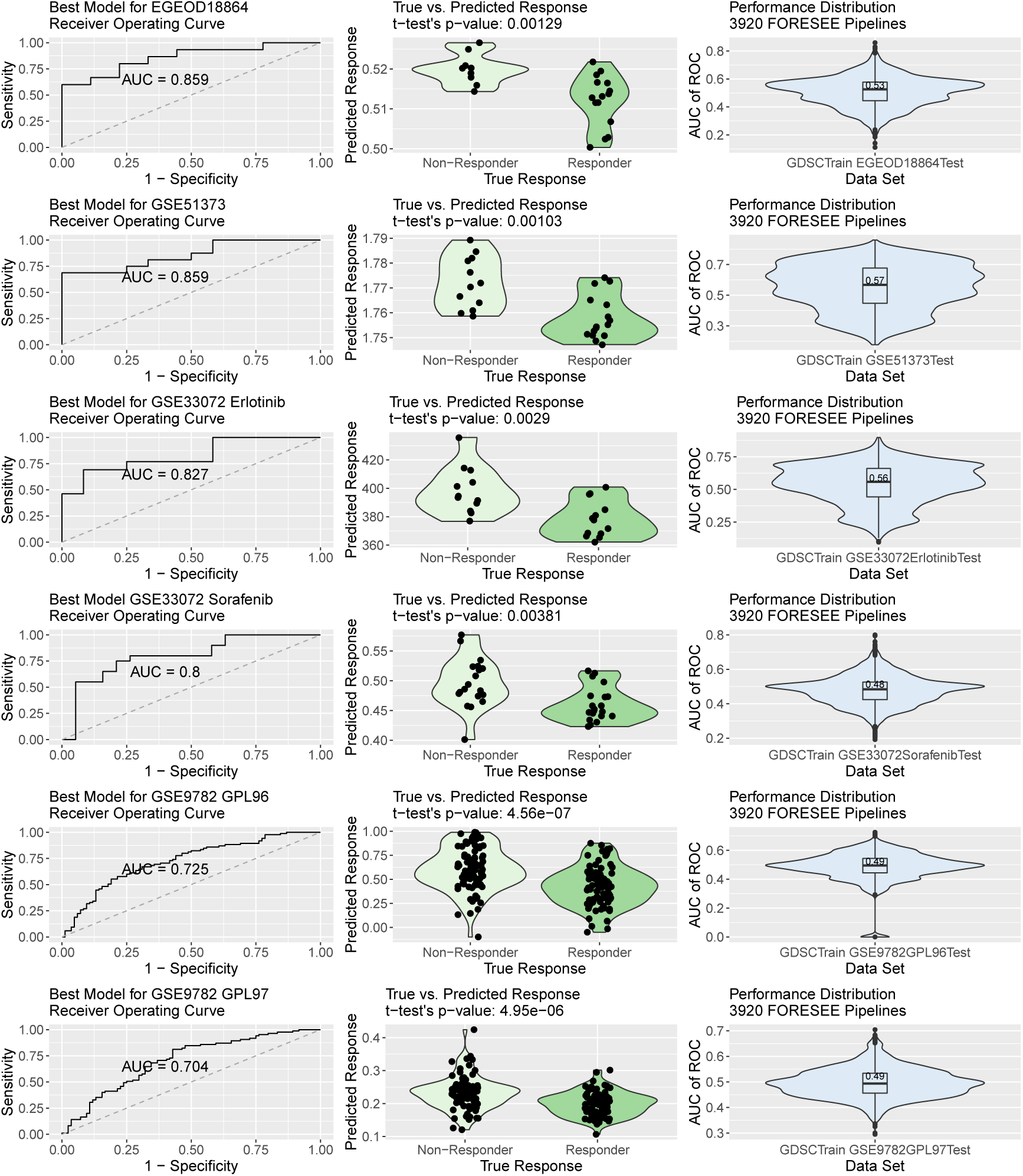
Portrayal of translational models that used the FORESEE package to train models on GDSC cell line data and subsequently predicted patient drug response of GSE18864, GSE51373, GSE33072Erlotinib, GSE33072Sorafenib, GSE9782GPL96 and GSE9782GPL97 patients. The settings for the respective best modeling pipelines can be found in Table 1. The patient responses were binarized as described in the paragraph Patient Response Preprocessing.

**Figure S2:**
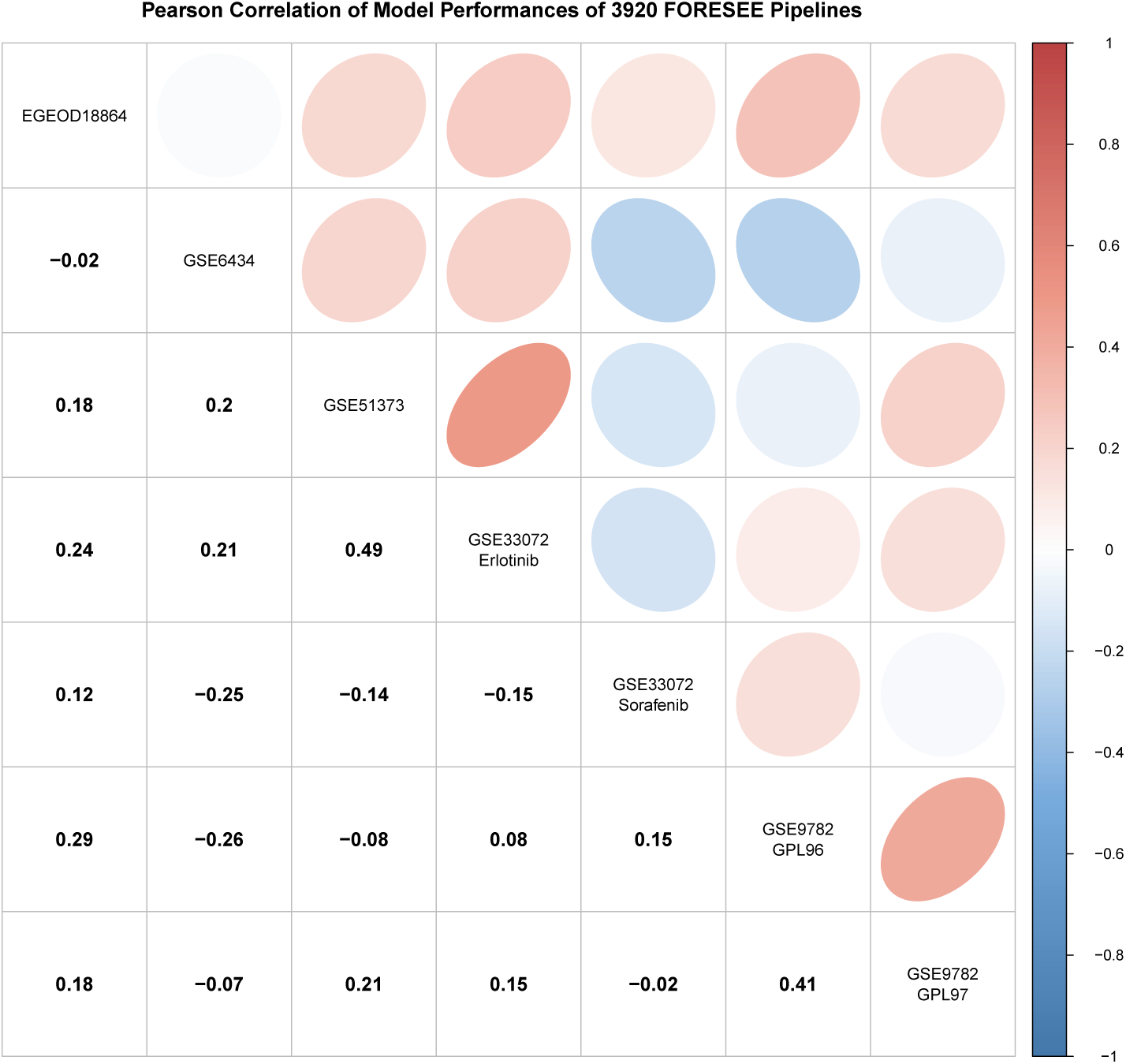
Pearson correlation of the performances of 3920 FORESEE pipelines for seven different patient data sets: GSE6434, EGEOD18864, GSE51373, GSE33072 Erlotinib cohort, GSE33072 Sorafenib cohort, GSE9782 GLP96 cohort and GSE9782 GLP97 cohort.

**Figure S3:**
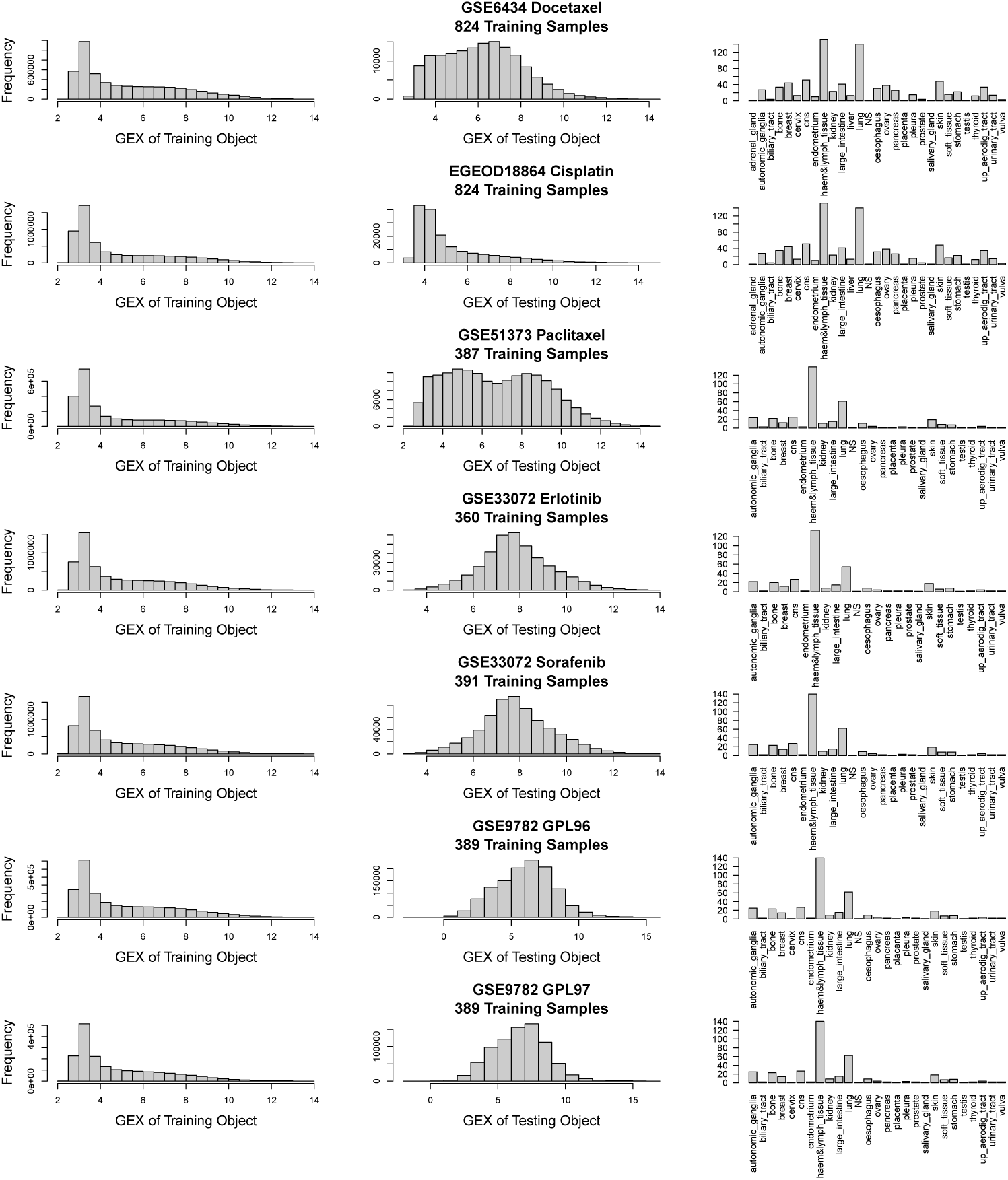
Distributions of the gene expression values of the cell line training data sets (A), the gene expression values of the patient testing data sets (B) and the cell line training data sets’ composition of tissues of origin (C) for seven different patient data sets: GSE6434, EGEOD18864, GSE51373, GSE33072 Erlotinib cohort, GSE33072 Sorafenib cohort, GSE9782 GLP96 cohort and GSE9782 GLP97 cohort.

